# The triose phosphate/phosphate translocator exports photosynthetic glyceraldehyde 3-phosphate from chloroplasts to trigger antimicrobial immunity in plants

**DOI:** 10.1101/2024.01.10.574840

**Authors:** Deng-Pan Zuo, Bin Wang, Yu-Zi Liu, Zheng-Song Chen, Ru-Jian Hu, Meng-Jun He, Zong-Ying Zhang, Ying Wang, Cheng-Gui Han

## Abstract

Chloroplasts play a crucial role in plant immunity against invading microbes. However, it remains poorly understood whether photosynthetic metabolites from chloroplasts participate directly in host defenses. Here, we uncovered *Arabidopsis thalinana* triose phosphate/phosphate translocator (AtTPT), a known translocator for chloroplast inner membrane, plays an indispensable role in suppressing virus infection and evoking defense responses. Interestingly, overexpression of AtTPT impairs virus accumulation in plants, while loss-of-function tpt3 mutants exhibit an increased viral load. The antiviral activity of AtTPT requires its phosphate transport capacity, implying that it actually functions through its metabolite(s). To this end, we found that glyceraldehyde 3-phosphate (GAP), one of AtTPT’s translocated metabolites, can drastically enhance expression of defense-related genes and prominently induce defense signaling pathways. More excitingly, AtTPT or GAP robustly restricts the proliferation of multiple types of phytopathogens. Collectively, we propose that AtTPT exports GAP to mediate broad-spectrum resistance to pathogens, which provides new insights into the mechanism underlying the chloroplast-mediated immunity by a photosynthetic metabolite.

## Introduction

Chloroplasts are organelles that carry out photosynthesis in the plant cell and responsible for the synthesis of phytohormone precursors, amino acids, carbohydrates, and lipids (*1, 2*). A large body of literature has documented core functions of chloroplasts in oxygenic photosynthesis and primary metabolism (*3*). Chloroplasts, as environmental sensors, respond to external stresses by generating chemical signals that regulate the expression of nuclear-encoded chloroplast genes (*4, 5*); they also participate in reprogramming the synthesis of primary and secondary metabolites to fine-tune phytohormone productions and intracellular communications via chloroplast-associated retrograde signaling (CRS) (*6–10*). Interestingly, beyond its above-mentioned roles in photosynthesis and signaling transduction, chloroplast also functions as a hub in immune responses and also as an important battlefield for attack and defense between plants and pathogens (*11–14*).

Metabolites from CRS pathways engage in multiple physiological and molecular processes in plant development and disease resistance, and these include tetrapyrroles, phosphoadenosines, carotenoid oxidation products, isoprenoid precursors, and hydrogen peroxide (*8, 10*). Additionally, CRS is of vital importance for chloroplast-mediated immunity. For instance, the chloroplast envelope forms tube-like extension, stromules, that convey NRIP1 (N receptor-interacting protein 1) and hydrogen peroxide (H_2_O_2_) to the nucleus, modulating CRS to reinforce ETI (Effector-Triggered Immunity) (*15, 16*). The SAL1 (nucleotidase/phosphatase)-PAP (3’-polyadenosine 5’-phosphate) retrograde signaling activates the SA (Salicylic acid) and JA (Jasmonic acid) signaling pathways, thereby enhancing plant immunity against the hemibiotrophic pathogen *Pseudomonas syringae* pv. tomato DC3000 (*Pst* DC3000) and the necrotrophic pathogen *Pectobacterium carotovorum* subsp. *carotororum* EC1 (*17*). MEcPP (2-C-methyl-d-erythritol-2,4-cyclopyrophosphate), one of the CRS metabolites, initiates a signaling cascade responding to both biotic (such as insects and viruses) and abiotic (like high light and chemical damages) stresses. MEcPP that accumulates in the cytosol boosts the systemic stress response (SSR) and accordingly, initiates the biosynthesis of defense-associated metabolites, such as pipecolic acid (Pip) and N-hydroxypiperic acid (NHP), leading to the production of SSR activators in response to these stresses (*18–20*).

Beyond governing redox and hormonal homeostasis, CRS also functions in regulating chloroplast-photosynthetic metabolite-linked pathways (*21*), but the underlying mechanism remain poorly understood. The chloroplast outer membrane forms an interfacial surface between the chloroplast and cytoplasm and is permeable to small molecules, while its inner membrane is less permeable and studded with specified translocators and ion channels for coordinating energy and metabolism between chloroplasts and the cytoplasm and nucleus (*4, 22, 23*). The triose-phosphate/phosphate translocator (TPT), one of the chloroplast-inner-membrane translocators, exports photosynthetic triose phosphates from chloroplasts and simultaneously imports inorganic phosphates from the cytosol (*24, 25*). TPT belongs to the plastid phosphate translocator (pPTs) family (*23*). TPT is an abundant protein in the cell, ensuring efficient carbon fixation in the chloroplast (*26*). The “rocker-switch” motion of the TPT molecule in substance translocation has been recently revealed by the crystal structure of TPT from the thermophilic red algae *Galdieria sulphonaria*, and its key residues for metabolites binding are found to be conserved evolutionarily and functionally in higher plants (*27, 28*). TPT is required for the short-term adaptation of plants responses to high light, involving apetala2/ERF transcription factors (key regulators for various stress responses) and the rapid phosphorylation of mitogen-activated kinase 6 (MPK6, translating external signals into cellular responses) (*21, 29*). Triose phosphates (TPs) including glyceraldehyde-3-phosphate (GAP) and dihydroxyacetone phosphate (DHAP), constitute the main products synthesized from photosynthetic carbon assimilation in chloroplasts and are thereafter exported to the cytoplasm mainly for sucrose synthesis (*30, 31*).

In chloroplasts, GAP, a photosynthetic metabolite, is synthesized by the Calvin-Benson cycle. Plants use Rubisco (ribulose-1,5-bisphosphate carboxylase/oxygenase) to incorporate one molecule of carbon dioxide into the second carbon of the five-carbon sugar, ribulose 1, 5-bisphosphate (RuBP). The resulting compound is extremely unstable and immediately breaks down into two molecules of 3-phosphoglycerate (3-PGA). 3-PGA is reduced by NADPH (produced in the photoreaction) with the consumption of ATP, producing glyceraldehyde 3-phosphate (GAP) (*32*). As an important intermediate for cellular processes and events, GAP is also involved in the pentose phosphate pathway as well as various other processes like glycolysis and gluconeogenesis (*33*).

Brassica yellows virus (BrYV) is a newly identified virus species mainly infecting crucifer crops with widespread distribution in Asia (*34–36*). BrYV belongs to the genus *Polerovirus* and the family of *Solemoviridae*. It is a phloem tissues-restricted and aphid-transmitted virus which spreads in a persistent, circulative, and non-propagative manner (*37–39*). BrYV genome comprises a positive-sense RNA of 5,666-5,678 nt with seven open reading frames (ORFs) encoding seven proteins P0, P1, P2, P3a, P3 (coat protein, abbreviated CP), P4 (movement proteins, MP), and RTP (a read-through protein fusing ORF3 and ORF5) (*34, 40*). The virus evades autophagy and proteasomal degradation by means of its P0 protein, a viral suppressor of RNA silencing (VSR), that interacts with plant S-phase kinase-associated protein 1 (SKP1) (*41*); P0 also facilitates viral pathogenicity by associating with host RAF2 (Rubisco Assembly Factor 2) and then impairing its antiviral function through altering its subcellular localization (*42*). The P3a protein aids the virus in long-distance movement (*43, 44*). BrYV infection or *Arabidopsis* plants harboring a transgene of *MP* significantly upregulate expression of anthocyanin biosynthesis genes and three sucrose-phosphate synthase genes, implying that the MP may interfere with host carbohydrate metabolism and distribution (*45*).

That plants modulate their endogenous carbohydrate metabolism and proteome appears to be a general defense strategy while fighting microbial infections (*46–48*). Triose phosphates exported by TPT are components in retrograde metabolic signaling regulating expression of nuclear-encoded photosynthesis genes. However, TPT or its exported metabolites have never been directly linked to disease resistance or defense signaling. More importantly, how chloroplasts contribute to defense against infections remain incompletely understood. Here, we identified the chloroplast inner membrane transporter, the triose phosphate/phosphate translocator (TPT), as a key regulator for plants to evoke antiviral responses, and this relies entirely on its metabolites transport capacity, which guided us to photosynthetic metabolites that TPT transports across the chloroplast membrane. Among TPT-transported metabolites, only GAP can effectively bring about host immune responses to virus infection. At the mechanistic level, GAP enhances expression of components in defense-related pathways and activates the MAPK signaling, both directly inhibiting virus spread. Excitingly, either overexpression of TPT or extra provision of GAP greatly promotes host disease resistance to multiple pathogenic-microbe species. Our findings provide new insights into the mechanism underlying the chloroplast-mediated immunity by discovering novel components in the CRS pathway and shed light on the involvement of chloroplast metabolism in disease resistance, potentially paving novel avenues for targeted intervention of microbes-caused plant diseases.

## Results

### TPT is required for *A. thaliana* plants to inhibit BrYV infection

The known role of TPT in plants is to export triose phosphates derived from CO_2_ assimilation in the light period and to support sucrose synthesis in the cytosol for supplying carbohydrates to the whole plant (*31*). To test whether TPT is involved in biotic stresses like viral infection. Col-0 and *tpt3* (a T-DNA insertion mutant of *tpt* (allele *tpt3*, SALK_093334) (*49*)) were inoculated with BrYV (transmitted by aphids) (*39*). In comparison to WT (Col-0), the abundance of BrYV CP, MP protein and RNA were significantly increased in virus-infected *tpt3* plants (Fig. 1, A and B). Furthermore, we generated transgenic *Arabidopsis* plants constitutively overexpressing *AtTPT-3xFlag* (AtTPT, GenBank: NM_001085251) under the control of the 35S promoter; this include *AtTPT* overexpression lines 3/4 (OE3/4) and genetic complementation lines 13/14 (COM13/14) in the background of Col-0 and *tpt3* respectively (Fig. 1C). Next, these transgenic plants were challenged with BrYV (transmitted via aphids). In comparison to WT (Col-0), *tpt3* plants developed more severe symptoms 25 dpi (Fig. 1D) concomitantly with an elevated virus load as represented by significantly increased MP and RNA abundance (1.6-fold and eight-fold respectively), whereas virus titers in COM13, COM14, OE3, and OE4 plants became appreciably restrained relative to *tpt3* (Fig. 1, E and F). Taken together, these results reveal the important role of TPT in suppression of viral infection.

**Fig. 1.**
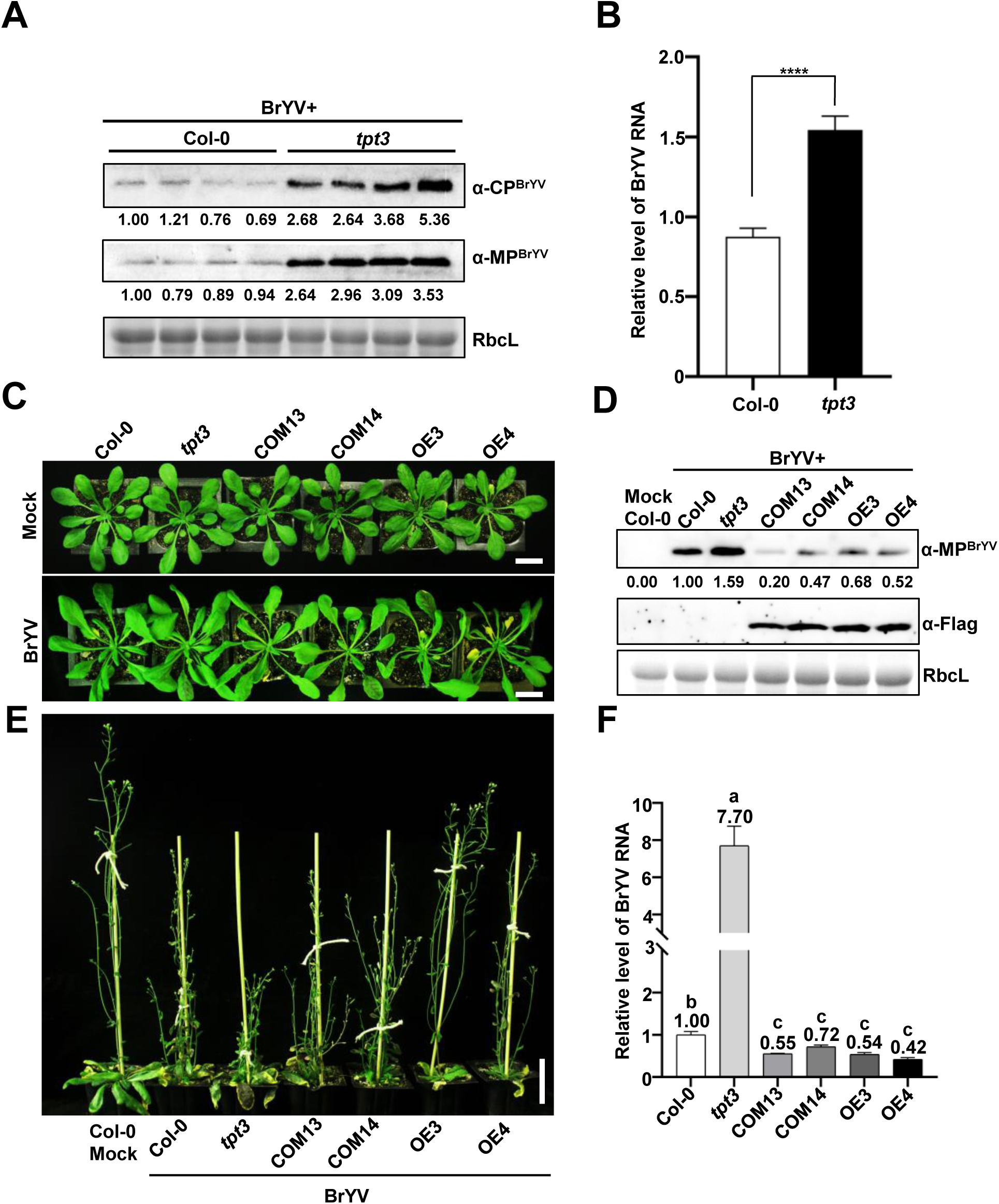
TPT is required for *A. thaliana* plants to inhibit BrYV infection. **(A)** Western blot analysis of BrYV’s CP and MP in BrYV systemically infected leaves of Col-0 and *tpt3* plants at 7 dpi. RbcL serves as a loading control.(**B**) RT-qPCR analysis of BrYV RNA accumulation in systemically infected leaves at 7 dpi. Error bars mean SD, and letters above the columns indicate significant differences (ANOVA, P<0.05). (**C**) Phenotypes of Col-0, *tpt3*, complementation lines (COM13 and COM14), and overexpression lines (OE3 and OE4) plants infected with or without BrYV (transmitted by aphids) at 14 dpi. Scale bar = 2 cm. (**D**) At 25 dpi, long-term systemic symptoms of Col-0, *tpt3*, COM13, COM14, OE3, and OE4 plants inoculated with BrYV. Scale bar = 5 cm. (**E**) Western blot analysis of BrYV’s MP in BrYV systemically infected leaves of Col-0, *tpt3*, COM13, COM14, OE3, and OE4 plants at 7 dpi. RbcL works as a loading control. (**F**) RT-qPCR analysis of BrYV RNA accumulation in systemically infected leaves at 7 dpi. Error bars mean SD, and letters above the columns indicate significant differences (ANOVA, P<0.05).

### AtTPT residues key for substance transport are needed for suppression of BrYV infection in *A. thaliana*

AtTPT belongs to the pPTs (plastid phosphate translocators) family and its key residues involved in metabolites binding are highly conserved, especially at H184, K203, Y336, K359, R360, and E206 (*27, 28*) (Fig. 2A). To further validate whether TPT’s transport activity is required for its antiviral function, we generated *AtTPT* mutants (in the *tpt3* background) harboring loss-of-function mutations at above-mentioned residues of AtTPT, and their expression is driven by the 35S promoter. Four independent *tpt*3 mutant lines bearing either H184A or R360A were selected to be challenged with BrYV, including TPT^H184A^OE/*tpt3*-1, TPT^H184A^OE/*tpt3*-8, TPT^R360A^OE/*tpt3*-21, and TPT^R360A^OE/*tpt3*-22 (see Materials and Methods for details). It has been reported that the *tpt* mutant deficient in metabolites transport favors starch biosynthesis in the chloroplast so the level of starch could serve as an indicator of the TPT activity (*50*).To validate the transport activity of AtTPT in the complement plants, starch contents were measured in Col-0, *tpt3*, COM13, COM14, TPT^H184A^OE/*tpt3*-1, TPT^H184A^OE/*tpt3*-8, TPT^R360A^OE/*tpt3*-21, and TPT^R360A^OE/*tpt3*-22. The genetic complement plants accumulated a higher level of starch than that in the Col-0 plants (Fig. 2B). The complement plants showed no abnormalities in growth and development relative to Col-0 and *tpt3* plants with or without infection by BrYV at 14 dpi (Fig. 2C). Western blotting and RT-qPCR analyses using leaves from systemically virus-infected plants revealed about 2-6-fold increase of viral CP, MP, and RNA in *tpt3* plants, as well as in TPT^H184A^OE/*tpt3*-1, TPT^H184A^OE/*tpt3*-8, TPT^R360A^OE/*tpt3*-21, and TPT^R360A^OE/*tpt3*-22 plants, as compared to the wild-type plants (Fig. 2, D and E). These results indicate that the transport activity of AtTPT is essential for its antiviral function.

**Fig. 2.**
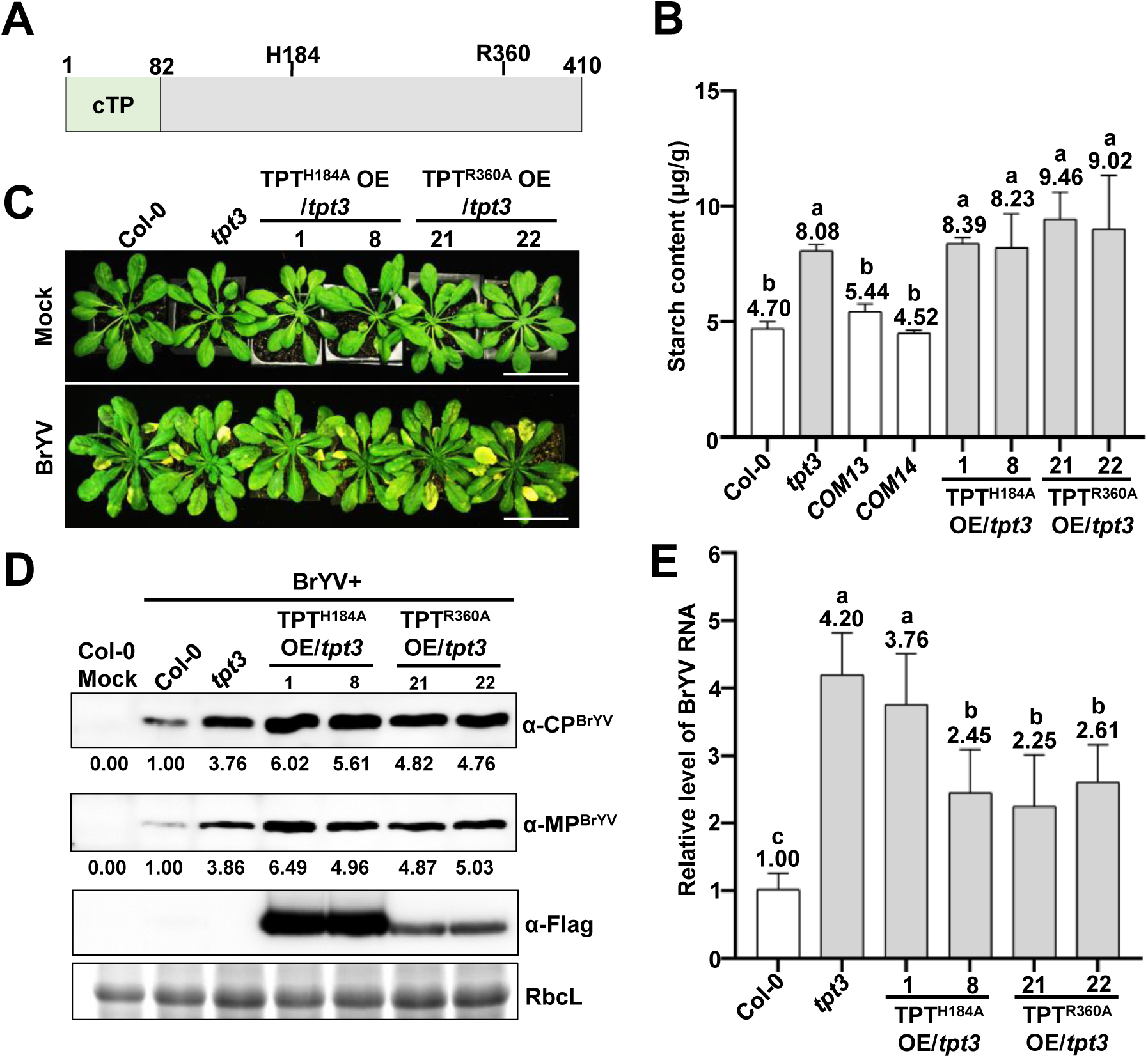
AtTPT residues key for substance transport are needed for suppression of BrYV infection in *A. thaliana*. (**A**) Schematic representation of AtTPT and its conserved amino acids required for phosphate transport (cTP: chloroplast transit peptide; H184, R360 are conserved residues for phosphate transport). (**B**) Starch-content measurement of Col-0, *tpt3*, COM13, COM14, TPT^H184A^OE/*tpt3*-1, TPT^H184A^OE/*tpt3*-8, TPT^R360A^OE/*tpt3*-21, and TPT^R360A^OE/*tpt3*-22.The asterisk (*) indicates significant differences in statistics (ANOVA, P<0.05). (**C**) Developmental and disease phenotypes at 14 dpi in the stated plants treated with mock or challenged with BrYV. Scale bar = 5 cm. (**D**) Western blot detection of BrYV CP and MP in systemically infected leaves at 7 dpi. (**E**) RT-qPCR analysis of BrYV RNA accumulation in systemically infected leaves at 7 dpi. Error bars mean SD, and letters above the columns represent statistically significant differences (ANOVA, P<0.05).

### AtTPT confers broad-spectrum resistance to multiple types of pathogens in *A.* ***thaliana* plants**

As TPT is conserved in all green plants especially for its amino acid residues recognizing its sugar substances among pPTs subtypes (*27*), we wonder if AtTPT could confer protection for plants against viruses from other families. We inoculated TuMV-GFP or CMV on Col-0 and *tpt3* plants and tracked the viruses in leaves by UV light and/or western blot analysis. The results showed that *tpt3* plants accumulated more TuMV-GFP (Fig. 3, A and B) and CMV (Fig. 3, C and D) than WT 14 dpi, demonstrating a critical role of AtTPT in virus suppression.

**Fig. 3.**
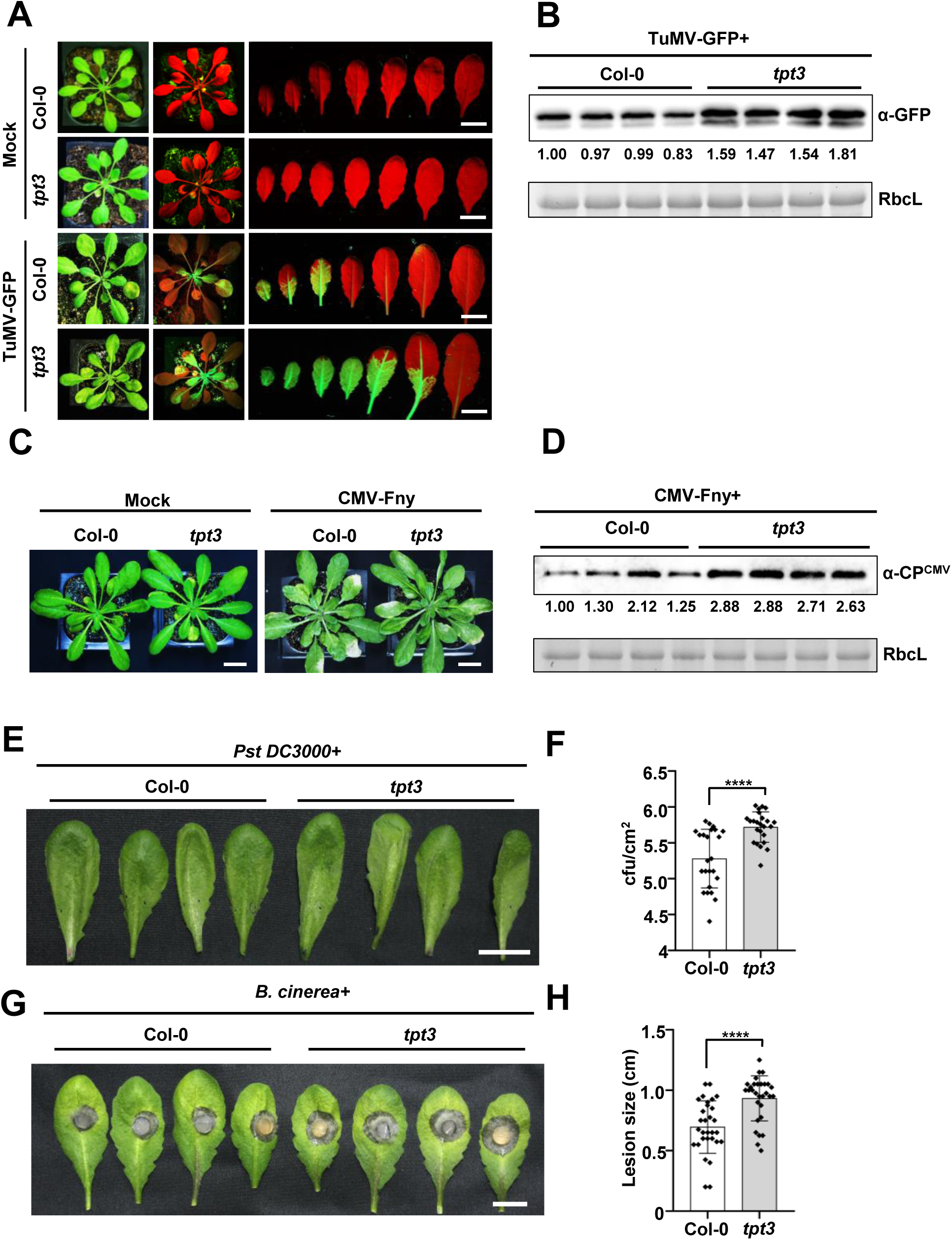
AtTPT confers broad-spectrum resistance to multiple types of pathogens in *A. thaliana* plants. (**A**) TuMV-GFP distribution at 9 dpi in Col-0 and *tpt3* plants that were infected with TuMV-GFP. Scale bar = 2 cm. (**B**) Western blotting showing TuMV-GFP levels in systemically infected leaves at 9 dpi. Values below the GFP blot was derived from densitometric analysis of the protein bands above. RbcL is a reference for equal loading. (**C**) Growth and disease symptoms at 10 dpi of Col-0 and *tpt3* plants that were either treated by a mock buffer or infected with CMV-Fny. Scale bar = 2 cm. (**D**) Western blotting showing CP^CMV^ in systemically infected leaves at 10 dpi. The level of CMV was represented by densitometric analysis of the CP^CMV^ bands, RbcL is used as a loading control. (**E**) Relative to Col-0, the *tpt3* plants displayed higher susceptibility to *Pst* DC3000. (**F**) Reproduction of *Pst* DC3000 in infiltrated leaves from Col-0 and *tpt3.* Colony-forming unit (cfu)/cm^2^ was the number of colonies per cm^2^ on the surface of agar medium (see Materials and Methods for details). Data are mean SDs (n = 24), and asterisks indicate a statistically significant difference. ****P< 0.0001 according to a Student’s t-test. (**G**) Representative images of *B. cinerea-*infected leaves that were detached from four-week-old Col-0 or *tpt3* plants. The images were taken at 2 dpi. The scale bar is 2 cm.(**H**) Comparison of lesion sizes in the Col-0 and *tpt3* plants at 2 dpi. Data are shown as the mean±SD with n=30, and asterisks indicate a statistically significant difference: ****P<0.0001 by the Student’s t-test.

As shown above, AtTPT can boost the innate immune system to combat viruses, which made us wonder whether they could also promote resistance to other types of pathogens. We chose *Pseudomonas syringae* pv. *tomato* DC3000 (*Pst* DC3000), a well-characterized bacterial pathogen infecting numerous plant species (*51*) and *Botrytis cinerea* (*B. cinerea*), a necrotrophic fungus that infects over 200 plant species (*52*) to challenge Col-0 and *tpt3* plants. In one set of experiments, rosette leaves of 4-week-old Col-0 and *tpt3* plants were infiltrated with *Pst* DC3000 and then collected to measure the bacterial load in the leaves 48 hpi (see Materials and Methods for details). Interestingly, *tpt3* mutants contained significantly more *Pst* DC3000 than that in the wild-type plants (Fig. 3, E and F). In the other set of parallel experiments but with *B. cinerea* as the pathogen, detached leaves from Col-0 and *tpt3* were challenged with the fungus, lesions appeared 3 dpi, and the average size of the lesions on the Col-0 leaves was 0.70 mm (n = 30) in diameter, whereas it became 0.93 mm (n = 30) on the *tpt3* leaves (Fig. 3, G and H). Altogether, these findings indicate that TPT in *A. thaliana* plants confers broad-spectrum resistance against pathogens.

### GAP mediates broad-spectrum resistance to multiple types of pathogens

Given that AtTPT requires residues essential in transport to restrict BrYV infection (Fig. 2), and TPT directly transports photosynthetic intermediates triose phosphates across the chloroplast envelope to the cytoplasm (see Introduction), we postulated that TPT triggers cellular immune responses via its translocated metabolites. We selected three candidate metabolites, GAP, DHAP, and 3-PGA (Fig. 4A), that have been reported to be mainly transported by TPT (*53*) and explored their effects on virus resistance. *N. benthamiana* is also a host of BrYV and it is more efficient and convenient to inoculate BrYV than *Arabidopsis* by agroinfiltration. Therefore, each of the selected metabolites (at 1 mM) was separately infiltrated with BrYV into *N. benthamiana* leaves. The treated leaves were collected for detecting the viral protein (MP) and RNA after two days. Interestingly, among the three, only GAP drastically lowered the viral RNA by ∼70%, as compared to the mock. The virus titer was unaffected by the DHAP or 3-PGA treatment (Fig. 4, B and C). To further characterize the dose of GAP with its antiviral effect, a titration assay using GAP at a series of concentrations of 1, 2.5, and 5 mM respectively was co-infiltrated with BrYV into *N. benthamiana* leaves. GAP displayed a dose-dependent effect on virus inhibition as supported by the reduced amount of viral protein (by 40-50, 70, and ∼90%, respectively) and RNA (by 30, 40, and 80%, respectively) (Fig. 4, D and E). The MP of BrYV was almost undetectable when the GAP concentration rose to 5 mM, indicating that a high dose of GAP can block viral propagation in plants. To confirm the antiviral activity of GAP in *A. thaliana* plants, the three metabolites (GAP, DHAP, and 3-PGA) were also evaluated for virus resistance using a transgenic *Arabidopsis* line that was transformed with the full-length infectious cDNA clone of BrYV (*45*) by spray application (every 12 hours for four times in total), similar to the results obtained from *N. benthamiana* (Fig. 4, B and C), only GAP displays a strong inhibitive effect on BrYV (by∼50% in comparison with the mock) (fig. S1, A and B). Altogether, the data indicate that AtTPT exerts its inhibitory effect on virus through GAP, and exogenous administration of GAP can inhibit virus infection in plants.

**Fig. 4.**
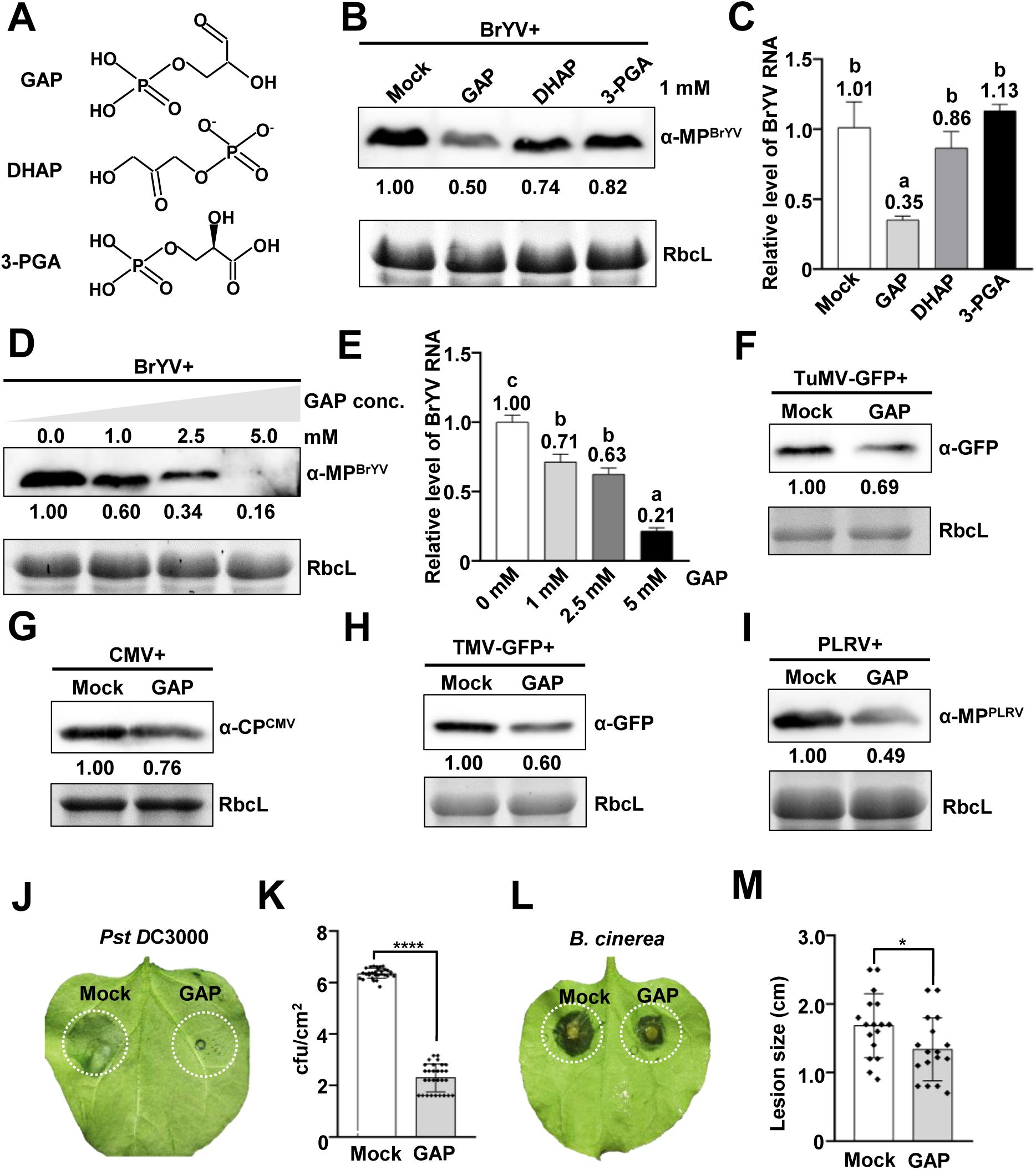
GAP confers broad-spectrum resistance to multiple types of pathogens. (**A**) Structural formula of glyceraldehyde-3-phosphate (GAP), dihydroxyacetone phosphate (DHAP), and glycerate 3-phosphate (3-PGA). (**B**) Effects of GAP, DHAP, or 3-PGA on virus accumulation in BrYV-inoculated *N. benthamiana* leaves. *N. benthamiana* leaves were infiltrated with BrYV as well as a mock buffer or 1 mM of GAP, DHAP, or 3-PGA and then harvested 48 hrs later for western blotting. (**C**) RT-qPCR analysis of BrYV RNA accumulation in inoculated leaves as treated in (C). Error bars indicate SD, and letters above show statist ical differences (P<0.05). (**D**) Western blot detection of BrYV’s MP using virus-inoculated leaves of *N. benthamiana* under the treatment of GAP at indicated concentrations (1.0∼5.0 mM). (**E**) RT-qPCR analysis of BrYV RNA accumulation in inoculated leaves as treated in (D). Error bars represent SD, and letters display statistical significance (ANOVA, P<0.05). (**F**) Western blotting showing that GAP (1 mM) inhibits TuMV-GFP accumulation in *N. benthamiana* leaves. The anti-GFP blot was used for detecting TuMV-GFP in western blotting. (**G**) Western blotting confirms that GAP (1 mM) lowers the CMV-GFP level in *N. benthamiana* leaves. The anti-CP^CMV^ blot was used for detecting CMV-GFP in western blotting. (**H**) Suppressive effects of GAP (1 mM) on TMV-GFP in *N. benthamiana* leaves. Anti-GFP was used for detecting TMV-GFP in western blotting. (**I**) Western blotting showing that GAP (1 mM) inhibits PLRV accumulation in *N. benthamiana* leaves. Anti-MP^PLRV^ was used for detecting PLRV in western blotting. (**J**) Necrotic symptoms at 2 dpi on *N. benthamiana* leaves caused by *Pst* DC3000 that was co-infiltrated with mock or GAP. Dashed outlines denote regions on the leaf infiltrated with mock or GAP. (**K**) Reproduction of *Pst* DC3000 in *N. benthamiana* leaves along with GAP or a mock buffer. Colony-forming unit (cfu)/cm2 was the number of colonies per cm^2^ on the surface of agar medium (see Materials and Methods for details). Data are mean SDs (n = 29), and asterisks indicate a statistically significant difference. ****P< 0.0001 according to a Student’s t-test. (**L**) Symptoms of necrosis on *N. benthamiana* leaves induced by *B. cinerea* in combination with 1 mM GAP or a mock buffer at 2 dpi. On the leaf surface, areas treated with mock or GAP were outlined. (**M**) Quantitative analysis of lesion sizes at 2 dpi from *N. benthamiana* leaves that were GAP-or sterile water-infiltrated and then inoculated with *B. cinerea*. Data are presented as the mean±SD (n = 17). *P< 0.1 based on a Student’s t-test.

In order to evaluate whether GAP can confer broad-spectrum resistance on virus infection, we infiltrated 1 mM GAP along with TuMV-GFP or CMV-GFP into *N. benthamiana* leaves. In line with the data acquired from BrYV (Fig. 4, B and C), GAP decreased accumulation of either TuMV-GFP or CMV-GFP in the leaves (Fig. 4, F and G). Similarly, we also noticed that GAP effectively restricts infections of TMV-GFP and PLRV in *N. benthamiana* leaves (Fig. 4, H and I). To test whether GAP can impact *Pst* DC3000 in *N. benthamiana*, leaves were inoculated with *Pst* DC3000 in the presence of GAP. Interestingly, *Pst* DC3000 infection was significantly obstructed in the leaves (Fig. 4, J and K). Compared to mock, GAP treatment clearly reduced the extent of *B. cinerea*-caused damages to the leaves (Fig. 4, L and M). All these findings indicate that GAP can mediate broad-spectrum protections for plants.

### GAP greatly induces defense-related genes expression in *N. benthamiana*

To better understand how GAP mediates immune responses mechanistically, we performed RNA-sequencing (RNA-seq) using *N. benthamiana* leaves infiltrated with or without 1 mM GAP. After 12, 24, and 48 hr of the GAP treatment, 15, 2,019, and 5,851 genes, respectively, were significantly differentially induced (Fig. 5A, Data S1). The KEGG (Kyoto Encyclopedia of Genes and Genomes) pathway enrichment analysis demonstrates that most of these over-represented genes in the transcriptomic data encode components functioning principally in the defensive network, including the hormone signaling, plant-pathogen interaction, the MAPK signaling, and starch/sucrose metabolism pathways (Fig. 5B). RT-qPCR was carried out to validate the expression of some of these defense-responsive genes: *NbPR1* (*N. benthamiana pathogenesis-related protein 1*) (*54*), *NbPR10* (*55*), *NbPDF1.2* (*N. benthamiana plant defensin 1.2*) (*56*), and *NbSABP2* (*N. benthamiana salicylic acid-binding protein 2*) (*57*). As expected, expression of all the tested genes appeared to be markedly increased by 24 and 48 h post GAP application: 2.9- and 13.3-fold for *NbPR1*, 1.9- and 7.7-fold for *NbPDF1.2*, 3.6- and 4.3-fold for *NbPR10*, and 28.3- and 79.6-fold for *NbSABP2* (Fig. 5, C to F), further confirming the RNA-seq result.

**Fig. 5.**
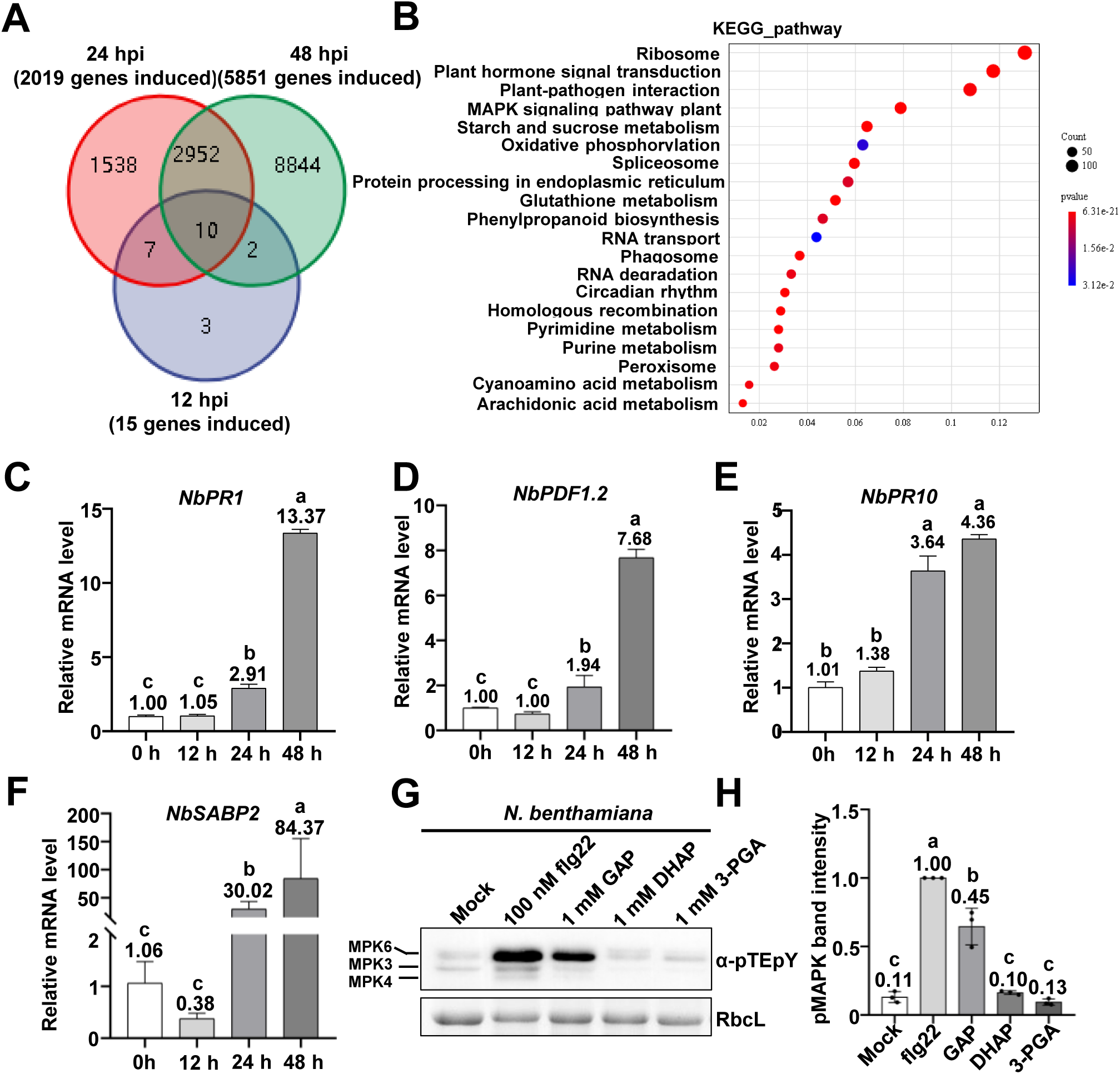
GAP induces expression of genes functioning in defense-related pathways in *N. benthamiana*. (**A**) A Venn diagram showing overlaps of upregulated genes in GAP-infiltrated leaves by 12, 24, and 48 hrs compared to the mock control. (**B**) Enrichment analysis of all upregulated genes from (A) by the Kyoto Encyclopedia of Genes and Genomes (KEGG) (https://www.genome.jp/kegg/). A color gradient lookup table indicates the p-value.(**C** to **F**) RT-qPCR analysis of *NbPR1*, *NbPDF1.2*, *NbPR10*, and *NbSABP2* expression in GAP-infiltrated *N. benthamiana* leaves. (**G**) Western blotting shows that only flg22 (a positive control) and GAP rather than DHAP and 3-PGA are capable of activating the MAPK pathway in *N. benthamiana* as represented by phosphorylated MAPK at Thr202/Tyr204 (detected by the pTEpY antibody). (**H**) Densitometric analysis of bands from the blot in (G) as well as its replicates (not shown here). Errors bars represent the SD of the mean (n = 3).

The MAPK signaling serves as a crucial pathway for turning on defensive genes expression and thereby boosting resistance to pathogens (*58*), including plant viruses (*59–61*). GAP is capable to suppress viral infection through inducing expression of immune-responsive genes (Fig. 4 and 5). This prompts us to investigate whether GAP can activate the MAPK signaling. To this end, we treated *N. benthamiana* plants with GAP, DHAP, or 3-PGA each at 1 mM for 15 min; flg22 (flagellin peptide 22 at the concentration of 100 nM (*61*)) was included in the test as a positive control, which carries a conserved epitope derived from bacterial flagellin and acts as a pathogen-associated molecular pattern (PAMP) in host-receptor recognition (*62*). Western blotting showed that only GAP, but not DHAP or 3-PGA modulates the MAPK signaling as fast and robustly as flg22 does (Fig. 6, G and H). This result and the transcriptome data (Fig. 5A) suggest that GAP first activates the MAPK signaling (within 15 min), and then the latter induces defense-related genes expression (∼24-48 h), although we cannot exclude the possibility that GAP or its derived metabolites directly promote transcription of defense genes. Collectively, our results indicate that exogenous application of GAP to plants can enhance defense-related genes expression and elicit MAPK signaling activation for hosts to fight infections.

**Fig. 6.**
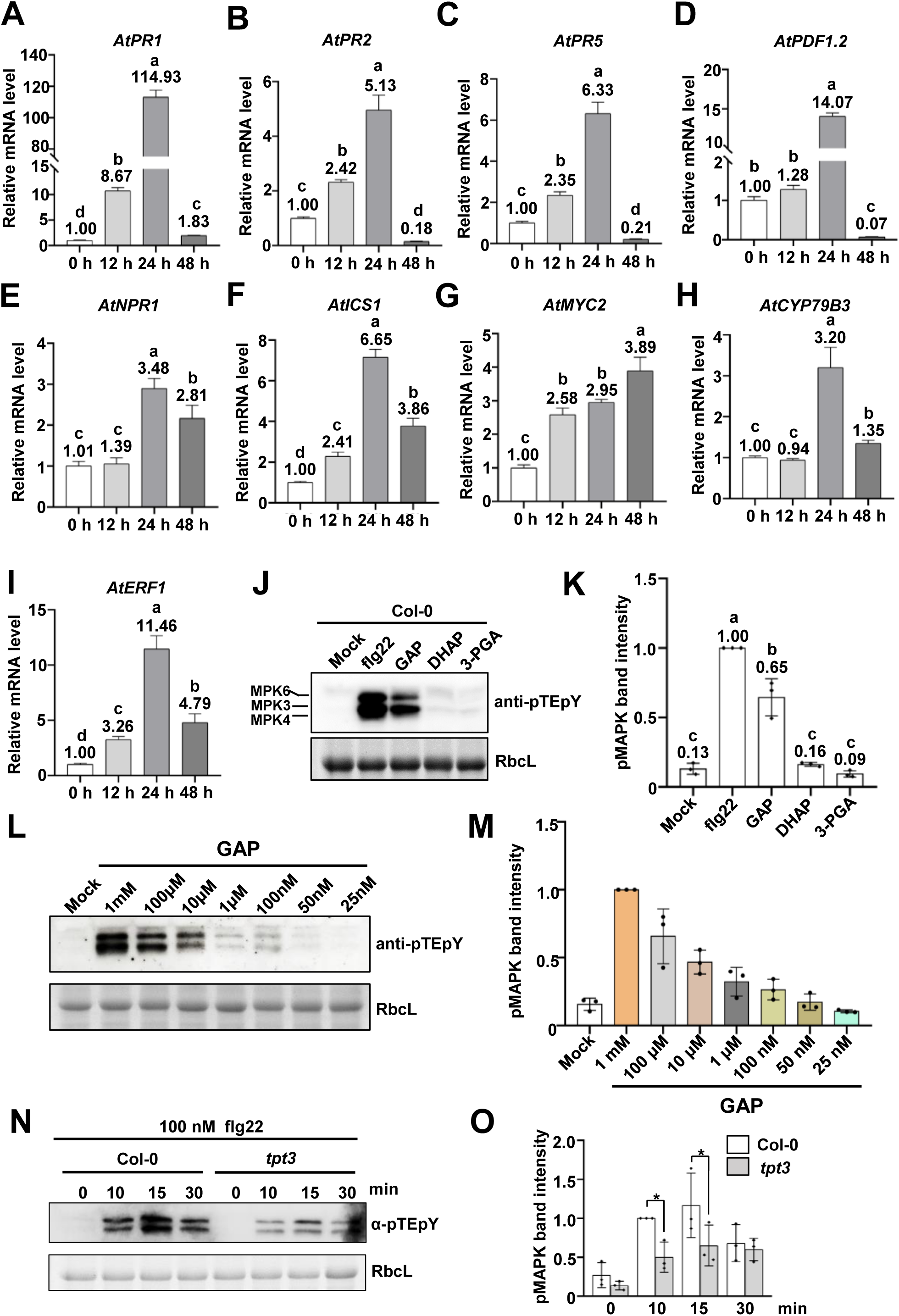
GAP triggers immune-related genes expression and MAPK signaling activation in *A. thaliana*. (**A** to **I**). RT-qPCR analysis of *AtPR1*, *AtPR2*, *AtPR5*, *AtPDF1.2*, *AtNPR1*, *AtICS1*, *AtMYC2*, *AtCYP79B3*, and *AtERF1* expression levels in GAP-sprayed *Arabidopsis* samples. Error bars and letters indicate SD and significant differences in statistics (ANOVA, P < 0.05) respectively. (**J**) Western blot show that, similar to flg22, GAP rather than DHAP or 3-PGA is able to elicit MAPK activation in the Col-0 and *tpt3* plants. RbcL is a loading control. (**K**) Quantitative measurements of bands from (J) and other replicates (data not shown). The asterisk (*) indicates significant differences in statistics (ANOVA, P < 0.05). (**L**) Western blot assays showed that flg22 activates the MAPK signaling less effectively in *tpt3* than in Col-0. RbcL works as a loading control. (**M**) Quantification of bands from (L) as well as other biological replicates (data not shown). Errors bars represent the SD of the mean (n = 3). (**N**) Western blot analysis showing the MAPK signaling activation in *Arabidopsis* leaves treated with GAP at indicated concentrations. (**O**) Densitometric analysis of bands from the blot in (a) as well as from other replicates (data not shown). Errors bars represent the SD of the mean (n = 3).

### GAP triggers immune-related genes expression and MAPK signaling activation in *A. thaliana*

The data presented above led us to an investigation of whether GAP could boost defense genes expression in *A. thaliana* as it does in *N. benthamiana* (Fig. 5). We sprayed GAP onto *Arabidopsis* leaves and collected them after 12, 24, and 48 h, respectively, to measure gene expression. Based on the transcriptome data from *N. benthamiana* leaves (Fig. 5), we selected *AtPR1* (*Pathogenesis-related Protein 1*, *PR1*) (*63*)*, AtPR2* (*64*)*, AtPR5* (*65*), and *AtPDF1.2* (*PLANT DEFENSIN TYPE 1.2*) (*66*), SA-related genes *AtNPR1* (*Non-expressor of pathogenesis-related genes 1*) (*67*) and *AtICS1* (*Isochorismate synthase 1*) (*68*), JA-responsive genes *AtMYC2* (*69*) and *AtCYP79B3* (*70*), and the ETHYLENE-responsive gene *AtERF1* (*71*) for RT-qPCR analysis. The result demonstrates that the GAP treatment dramatically enhanced expression of all the genes tested after 12 to 48 h (Fig. 6, A to I), in line with their expression profiles from *N. benthamiana* leaves (Fig. 5). Unlike in *N. benthamiana* (Fig. 5), GAP-induced transcription of multiple genes in *Arabidopsis* started to decline by 48 h, such as *AtPR1* and *AtPDF1.2* relative to 24 h (Fig. 6, A and D), which might be due to different metabolic rates of GAP in the two plant species. In addition, *Arabidopsis* plants were treated with 1mM of GAP, DHAP, or 3-PGA for 24 h, and induction of *AtPR1* and *AtPDF1.2* confirmed that GAP specifically promotes expression of defense-related genes in *A. thaliana* (fig. S2, A and B).

Subsequently, we tested effects of GAP, DHAP, or 3-PGA on MAPK activation in Col-0 plants. Leaves from Col-0 plants was treated with 100 nM flg22 peptide, 1 mM GAP, 1 mM DHAP, or 1 mM 3-PGA for 15 min. Similar to the positive control, flg22, GAP effectively triggers MAPK activation, whereas DHAP or 3-PGA entirely failed to do so (Fig. 6, J and K). The *in vivo* concentration of GAP in Col-0 plants was reported previously (*72*). To determine the minimal dose of GAP effective for MAPK activation, detached *Arabidopsis* leave discs were treated with GAP at 1 mM, 100 μM, 10 μM, 1μM, 100 nM, 50 nM, and 25 nM, respectively, and phosphorylated MAPK was followed by western blotting. The results showed that exogenous application of GAP at 50 nM or above can activate the MAPK signaling (Fig. 6, L and M). Taken together, external application of GAP can strongly induce expression of defense genes in *Arabidopsis*, including those controlling the synthesis of defense-related hormones.

The results above (Fig. 5G, and Fig. 6J) demonstrated that GAP behaves as a potential activator of the MAPK signaling. We wondered whether its transporter TPT is required to evoke the pathway. To address the question, leaf discs were excised from Col-0 and *tpt3* plants and treated with flg22 for 0, 10, 15, and 30 min. Western blot analysis showed that with respect to the WT, the activation of MPKs3/6 was remarkably attenuated in the *tpt3* plants that were exposed to flg22 (Fig. 6, N and O), suggesting that TPT functions as a critical, upstream component of the MAPK pathway. These results further confirmed that TPT and its transported GAP are involved in the MAPK signaling activation.

## Discussion

Rapidly growing literature reveals the crucial role of chloroplasts in establishing the host immunity to defend against invading pathogens. However, from the perspective of pathogenic microbes, completely destroying chloroplast functions would severely impair their propagation in the host due to their stringent reliance on hosts to obtain nutrients. But leaving chloroplasts fully intact would render infections almost impossible to achieve because of the active participation of chloroplasts in immune responses—a dilemma encountered by obligate parasites, such as plant viruses. Our findings are summarized in Fig. 7: TPT exports photosynthetic GAP to the cytosol to activate the defense signaling including MAPK as well as other unidentified pathways, which induces nuclear genes expression to promote disease resistance. We establish a molecular framework for the TPT- and GAP-mediated defense mechanism, exemplifying how a chloroplast protein and its cargo metabolite engage in host defense to pathogens.

**Fig. 7.**
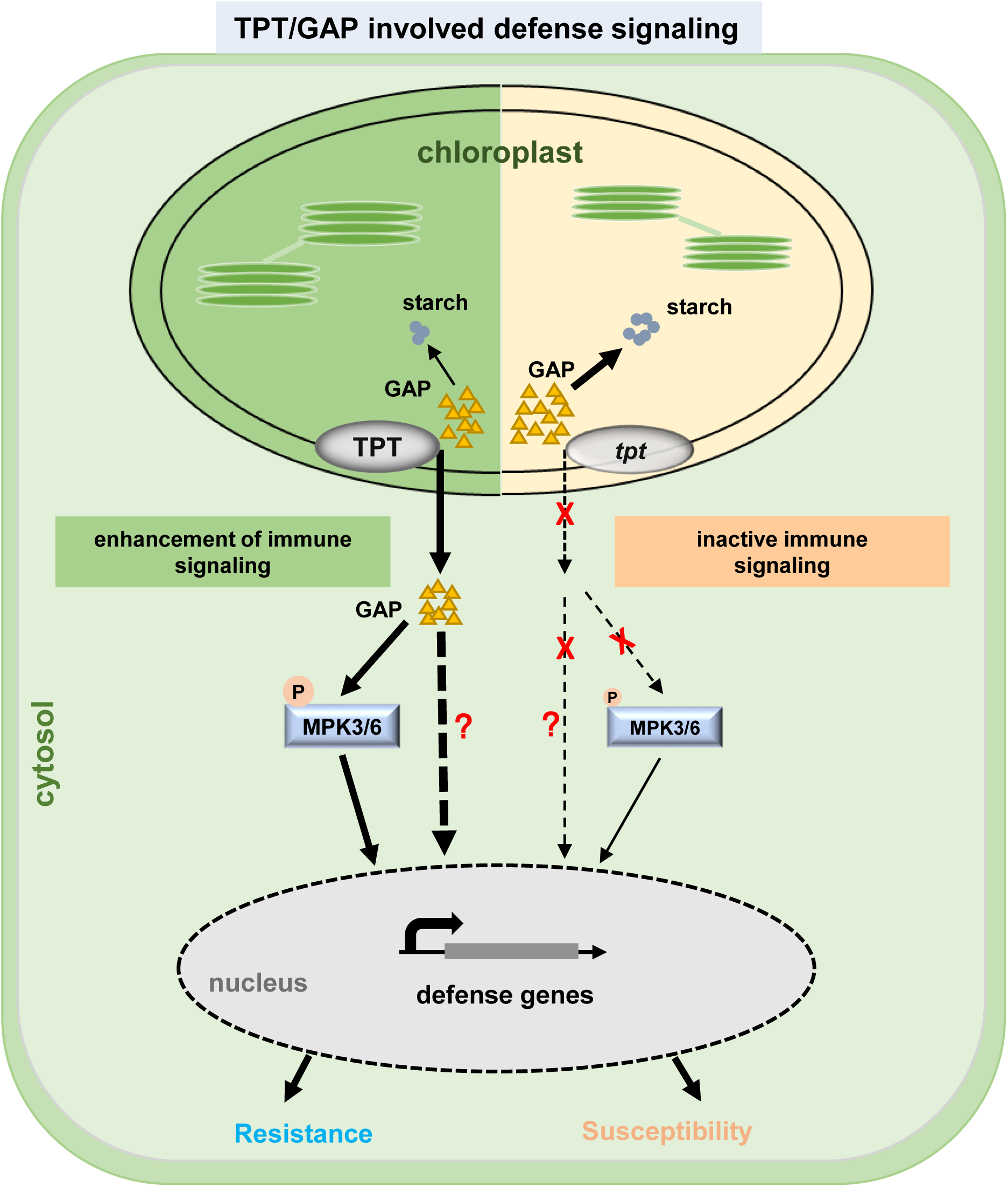
Working model for TPT/GAP-activated defense pathways in plants. TPT functions in chloroplasts inner membrane to translocate triose phosphate to the cytosol. GAP that is translocated by TPT from chloroplasts enhances expression of genes mediating defensive responses in normal conditions; But in *tpt* mutant plants, the amounts of GAP severely droped in the cytosol which makes immune-responsive pathways blunt, such as the MAPK signaling and defense-genes induction, thereby making plants more susceptible to pathogens. Thicker arrows mean stronger effects or higher efficiency, the dash arrows with a question mark imply uncertainties, and the larger phosphorylation symbol on MPK3/6 represents phosphorylation to a greater extent.

Chloroplast inner membrane serves as the major interface controlling the distribution of photo-assimilates between the chloroplast and cytosol and thereby determining plant growth, development, and metabolic balance (*73*). TPT is a crucial transporter on the chloroplast inner membrane, as substantiated by the observation that a *tpt-*deletion mutant of *Arabidopsis* contains more starch than that in wild-type in the light (*50*). In the retrograde signaling, TPT exports metabolites (TPs) and subsequently activates MPK6 to regulate expression of nuclear AP2/ERF-TFs (apetala2/ethylene response factor transcription factors) like ERF6, RRTF1, ERF104, and ERF105 for coping with the high light stress (*21*). However, it was unknown that which substance metabolite of TPT mediates MPK6 activation, and whether TPT and its transported TPs participate directly in responses to biotic stresses. As AtTPT inhibits virus infection and this relies on its transport activity (Fig. 1, and 2), we examined effects of metabolites known to be transported by AtTPT on virus suppression. Interestingly, only GAP can act potently in constraining virus infection (Fig. 4). Moreover, GAP can specifically induce defense genes expression and can also activate the MAPK signaling (fig. S2, A and B, and Fig. 5G, and Fig. 6, J and L). These observations shed light on the complexity underlying chloroplast-associated retrograde signaling (CRS) in immune system activation and again highlight the coordination of chloroplast metabolism with the immune network. In this study, we also found that flg22-induced MAPK signaling was impaired in the *tpt3* mutants (Fig. 6, L and M), demonstrating the indispensable role of TPT in the MAPK signaling activation which confers TPT to mediate broad-spectrum resistance to pathogens.

Mechanistically, TPT overexpression or GAP administration enhances plant immunity through activating defensive signaling and inducing the immune transcriptional network (Fig. 5 and 6), evoking systematic defenses against a wide range of pathogens (Fig. 3 and 4). How GAP in the cytosol activates the immune signaling and drives the transcriptional network (Fig. 5 and 6) remains unclear and will be the subject of future genetic and biochemical studies. It might directly bind to and allosterically activate essential components in the immune signaling, such as certain transcription factors and kinases; an alternative possibility exists that metabolites derived from GAP instead of GAP itself might be the functional substance triggering immune responses. Future investigations of characterizing GAP-derived metabolites in disease resistance may be revealing.

In plant cells, GAP as an intermediate metabolite takes part in multiple carbohydrate involved pathways nucleating the carbon metabolism network. For example, GAP and DHAP are interconvertible constitutional isomers that are catalyzed by the triose phosphate isomerase (TPI) in the cytoplasm (*74*), and they are utilized to produce fructose 1, 6-diphosphate (F1,6BP) for sucrose synthesis. Alternatively, in glycolysis the glyceraldehyde-3-P dehydrogenase (GAPC) converts GAP to 1,3-bisphosphoglyceric acid, and then the latter is transformed to 3-PGA by the 3-phosphoglycerate kinase (PGK) (*75, 76*). GAPCs have consequential roles in autophagy, hypersensitive responses, and host innate immunity (*77, 78*). Silencing *GAPC* genes enhances *N*-gene-mediated resistance against TMV and pathogenic bacterium *Pseudomonas syringae* (*78*). Recently, it has been shown that CP19K encoded by RNA2 of CWMV (Chinese wheat mosaic virus, genus *Furovirus*) targets ATG3 and GAPC to form the CP19K-GAPC-ATG3 complex and upregulates GAPC expression to inhibit autophagy, which promotes CWMV infection (*79*). These findings agree with a model wherein the reduction of GAPC results in a higher GAP level, which promotes the antimicrobial immunity. Whether the underlying mechanism of GAP-mediated immune responses is related to autophagy still needs further evidence to prove. The host *OsTPI1.1* (*Oryza sativa* TPI 1.1) gene has been reported to function as a positive regulator in *XA3/XA26* (*80*)-mediated defense against *Xanthomonas oryzae* pv *oryzae* (Xoo), but *in vitro* and *in vivo* results are conflicting regarding its activity in controlling the ratio of GAP to DHAP. GAP has never been explicitly linked to disease resistance (*81*).

The major end products from photosynthesis consist of sugars such as glucose, sucrose, fructose, and trehalose and may have potentials to modulate expression of nuclear-encoded photosynthesis genes (*82–85*). It is surprising that GAP as a photosynthetic metabolite originated from chloroplasts can augment expression of nuclear genes especially those functioning in the defense network. In contrast, structurally similar metabolites DHAP and 3-PGA do not play appreciable roles in virus resistance (Fig. 4). The transcriptomic data reveal that GAP-inducible genes mainly include those involved in hormone signal transduction, plant-pathogen interaction, MAPK signaling pathway, and starch/sucrose metabolism (Fig. 5B), which contribute directly or indirectly to immune responses. GAP boosts expression of defense-related genes in *N. benthamiana* leaves (Fig. 5) and *Arabidopsis* (Fig. 6, A to I). For example, spraying GAP onto *Arabidopsis* leaves enhanced expression of genes required for the synthesis of classic defense hormones, such as SA, JA, and ET (Fig. 6, A to I). Based on the transcriptomic data and corresponding RT-qPCR validations, the GAP treatment robustly enhances expression of genes whose gene products work for defense-hormone syntheses and immune signaling. The SA, JA, ET, and MAPK pathways are intimately intertwined in the immune system (*86–88*). Thus, the hierarchy of components that govern the GAP-induced defensive network merits further investigations.

## Materials and Methods

### Plasmid construction

To make *AtTPT* mutants for *Arabidopsis* transformation, the *AtTPT* gene was first amplified by PCR and then introduced into the pMD19-T vector (TaKaRa, Dalian, China) to construct pMD19-T-AtTPT; inverse-PCR with pMD19-T-AtTPT as the template was performed to generate *AtTPT* mutants bearing substitutions to the key residues in substance binding—pMD19-T-AtTPTm (H184A, K203A, E206A, Y336A, K359A, or R360A). Subsequently, DNA fragments containing these mutations were cloned into the pMDC32-3xFlag (*89*) vector, resulting in pMDC32-AtTPT^H184A^-3xFlag, pMDC32-AtTPT^K203A^-3xFlag, pMDC32-AtTPT^E206A^-3xFlag, pMDC32-AtTPT^Y336A^-3xFlag, pMDC32-AtTPT^K359A^-3xFlag, and pMDC32-AtTPT^R360A^-3xFlag respectively. Infectious cDNA clones of BrYV, PLRV (*90*), TMV-GFP (*91*), CMV-GFP (*92*), TuMV-GFP (*93*), and CMV-Fny (*94*) infectious cDNA clones used in this study were previously described. Primers used for cloning and other molecular manipulations were listed in table S1. The sequences of plasmids used in this study were verified by Sanger sequencing (Tsingke Biotechnology).

### Plant materials and growth conditions

Transgenic plants for the purpose of overexpression are all in the Col-0 (Wild-type *A. thaliana*) background. pMDC32-AtTPT-3xFlag was transformed into the Col-0 plants to obtain transgenic lines of OE3/4. The T-DNA insertion line of *tpt3* (SALK_093334) was provided by Professor Shu-Hua Yang (College of Biological Sciences, China Agricultural University). Homozygous individuals were genotyped by PCR using gene-specific primers listed in table S1. Genetic complementation plants were generated by transforming WT or mutant AtTPT (H184A, K203A, E206A, Y336A, K359A, or R360A) (all cloned into pMDC32 respectively and each with a 3xFlag) into *tpt3* plants; the resultant lines for further investigations include COM13/14 (WT AtTPT-3xFlag), TPT^H184A^OE/*tpt3*-1/8 (AtTPT^H184A^-3xFlag), and TPT^R360A^OE/*tpt3*-21/22 (AtTPT^R360A^-3xFlag). *Arabidopsis* plants were grown in a chamber at 20-22°C with a 10-h-light/14-h-dark regime, while *N. benthamiana* plants were cultured at 24°C with a 16-h light/8-h dark cycle.

### Western blots

Western blotting was done with corresponding antibodies against GFP (1:5,000, EASYBIO, Cat. No. BE2002) and Flag (1:5,000, EASYBIO Cat. No. BE2005) respectively. Antibodies against CP^BrYV^, MP^BrYV^, MP^PLRV^, CP^CMV^, and CP^TMV^ were all homemade and reported previously. Phosphorylated MAPK was detected using the phospho-p44/42 MAPK antibody (for monitoring phosphorylated MAPK at Thr202/Tyr204 and marked as pTEpY in figures) (1:2,000 dilution, Cell Signaling Technology, Cat. No. 4370).

### Virus inoculation and detection

For inoculation assays with *N. benthamiana* leaves, infectious cDNA clones of indicated viruses were transformed into *Agrobacterium C58C1* (BrYV, PLRV) or *GV3101* (TuMV-GFP, CMV-GFP, and TMV-GFP), and then the latter was used to infiltrate *N. benthamiana* plants that have grown in size by about 3-4 weeks. Total protein and RNA were extracted from treated areas on the leaves, and viral proteins including CP and MP and RNA as representatives of virus abundance were analyzed by western blotting and RT-qPCR respectively. The detailed method for aphid-mediated transmission of BrYV was described previously (*39*). For TuMV-GFP and CMV infection assays, four-week-old *Arabidopsis* seedlings were inoculated mechanically with sap from TuMV-GFP-infected *N. benthamiana* leaves (ground in phosphate buffer [pH7.4]) or purified CMV virions (diluted to 5 ng/μL) as previously described (*94*). Fluorescence emitted from TuMV-GFP-infected *Arabidopsis* leaves was monitored under a long-wavelength UV lamp (Analytik Jena [former UVP], CA, USA) and recorded by a digital camera 14 dpi. Protein detection by western blotting was described in the section of “Western blots”.

### RNA extraction and RT-qPCR

Total RNA was extracted using the TRIzol reagent (Invitrogen, CA, USA) according to the manufacturer’s protocol. 2 μg of purified total RNA was treated with RNase-free DNaseI (Takara, Dalian, China) to remove contaminated genomic DNA. The first-strand cDNA was synthesized by the M-MLV reverse transcriptase (Promega, Madison, USA). RT-qPCR assays were performed with the GoTaq qPCR Master Mix (Promega, Madison, USA). Two genes, *AtActin2* for *Arabidopsis* samples and *NbEF1A* for *N. benthamiana* plants, served as references for normalization. All primer pairs used in RT-qPCR analysis were listed in table S1.

### RNA-Seq analysis

The *N. benthamiana* leaf samples were collected at 12, 24, and 48 h after mock or GAP treatment. Four leaves from one plant were pooled as one biological replicate, and each treatment included three biological replicates. Total RNA was extracted using the TRIzol reagent (Invitrogen, CA, USA). For each sample, 1 μg of purified total RNA was taken as the “input”. Sequencing libraries were constructed using the NEBNext Ultra RNA Library Prep Kit for Illumina (NEB, USA) following the manufacturer’s recommendations, and index codes were added to attribute sequences to each sample. Fragments in the library were purified with the AMPure XP system in order to select cDNA preferentially at 240 bp in length (Beckman Coulter, Beverly, USA). The clustering of the index-coded samples was performed via the cBot cluster generation system using TruSeq PE Cluster Kit v4-cBot-HS (Illumia) according to the manufacturer’s instructions. The prepared libraries were sequenced on an Illumina platform, generating paired-end reads. The raw reads were further processed with a bioinformatic tool, the BMKCloud online platform (www.biocloud.net). Hisat2 was used to map the read sequences to the reference genome. The efficiency of read alignments to the reference genome in each sample ranges from 95.11 to 95.80% as determined by detailed comparisons. Differential expression between two conditions/groups was determined by DESeq2, providing statistical analysis for assessing quantitative differences between samples based on the negative binomial distribution. The resultant p-values were adjusted with the Benjamini and Hochberg Procedure to control the false discovery rate, and genes with an adjusted p-value < 0.01 (by DESeq2) were regarded as differentially expressed between samples.

### Measurements of MAPK activation

Leaves from three- to four-week-old *Arabidopsis* or five-week-old *N. benthamiana* were soaked in sterile water overnight and then treated with 100 nM flg22 (GlpBio, Cat. No. GC32202) for 15 min; similarly, *Arabidopsis* or *N. benthamiana* leaves were handled with 1 mM GAP (GlpBio, Cat. No. GC43490), 1 mM DHAP (Sigma, Cat. No. D7137), or 1 mM PGA (Shanghai Yuanye Bio-Technology, Cat. No. S25079) for 15 min. Total protein was extracted using an extraction buffer (100 mM Tris [pH 6.8], 4% [w/v] SDS, 20% [v/v] glycerol, and 0.2% [w/v] bromophenol blue). Antibody against phosphorylated MAPK for western blotting was described under the section of “Western blots”.

### Quantification of starch content in plant leaves

Amounts of starch content in plants were measured with a kit (Suzhou Comin Biotechnology) in accordance with the manufacturer’s instruction. 1 g tissue from each sample was taken for quantification.

### Infections with *Pseudomonas syringae* and *Botrytis cinerea*

For *Pseudomonas syringae* (*Pst* DC3000), inoculation was done as previously described (*51, 95*) with minor modifications. Briefly, the *Pst* DC3000 strain was cultured at 28°C overnight and then diluted to a density at OD_600_ ∼0.0005. Four-week-old *Arabidopsis* plants were inoculated with a needleless syringe containing the diluted *Pst* DC3000 and then covered with a transparent plastic dome for 2 days for humidity maintenance. After taking photos of the whole plants, two inoculated leaves were collected, and 2 discs from each leaf were removed with a perforator (5 mm in diameter) and stored as one biological replicate. The leaf discs were ground up and diluted with sterile water. Bacterial suspension from ground leaf discs was plated on Luria–Bertani (LB) agar medium containing rifampicin (50 mg/μL), and the number of colonies grown on the plate was counted after 48 hrs of incubation at 28°C and divided by leaf surface areas to acquire values of colony-forming unit (cfu)/cm^2^. For GAP-involved inoculation assays, *Pst* DC3000 suspension was co-inoculated with 1 mM GAP or sterile water (mock) onto *N. benthamiana* leaves for 2 days. Photography, leaf discs removal, and colony counting were done in the same way as for *Arabidopsis*.

*Botrytis cinerea* (*B. cinerea*) infection was carried out as previously described (*52*). In brief, *B. cinerea* was cultured on potato dextrose agar medium at 28°C for 5-7 days. Agar plugs (2 mm in diameter) were used to inoculate detached leaves from five-week-old *Arabidopsis* plants. Images of the infected leaves were taken and stored, and the size of lesions was measured 2-3 dpi. For inoculation assays involving GAP, *N. benthamiana* leaves were pre-infiltration with 1 mM GAP or sterile water (mock); 24 hrs later, agar plugs (2 mm in diameter) containing *B. cinerea* were inoculated onto the pretreated areas for 2 days. Images of treated leaves were taken/stored, and the size of lesions was scored.

### Supplemental Data Sets

**table S1.** Primers used in this study.

### Data availability

The RNA-seq raw data generated in this study have been deposited into the NCBI database with the accession code: PRJNA896131, and the source data are provided with this paper.

## Acknowledgements

We thank Prof. Shu-Hua Yang for the *tpt3* mutant. We thank Drs. Jia-Lin Yu, Da-Wei Li, Xian-Bing Wang, and Yong-Liang Zhang at China Agricultural University for their valuable suggestions for this work. This work was supported by the National Natural Science Foundation of China to Cheng-Gui Han (32272494 and 31972240) and National Key R&D Program of China (No. 2023YFD1400300).

## Author contributions

CG.H. and DP.Z. conceived the study and designed the experiments. DP.Z. performed the experiments with the help of YZ.L., ZS.C., RJ.H., and MJ.H. Y.W. and ZY.Z. provided suggestions for this work. DP.Z., B.W., and CG.H. wrote and revised the manuscript. All authors analyzed the data.

## Competing interests

The authors declare no competing interests.

## Notes

### Competing Interest Statement

The authors have declared no competing interest.

