## Supplemental files for "The triose phosphate/phosphate translocator exports photosynthetic glyceraldehyde 3-phosphate from chloroplasts to trigger antimicrobial immunity in plants"

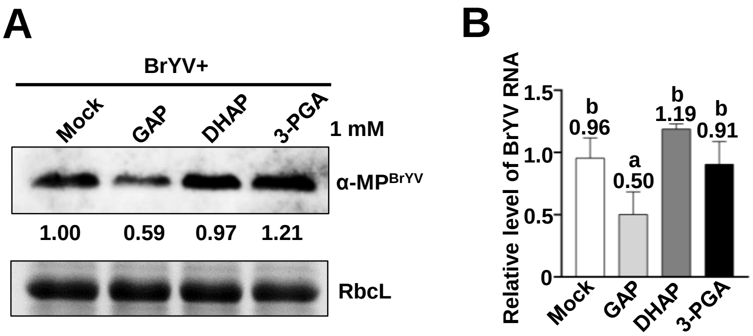


**Fig. S1. Glyceraldehyde-3-phosphate (GAP) restricts BrYV infection in *Arabidopsis thaliana*.** (**A**) The repressive effect of GAP on virus in transgenic plants carrying the BrYV genome. GAP, DHAP, or 3-PGA each at 1 mM was sprayed onto the leaves every twelve hours for four times in total, and western blot analysis was carried out for measuring BrYV abundance. (**B**) RT-qPCR analysis of BrYV RNA accumulation in systemically infected leaves that were treated by GAP, DHAP, 3-PGA, or mock. Letters represent significant differences in statistics (ANOVA, P<0.05).

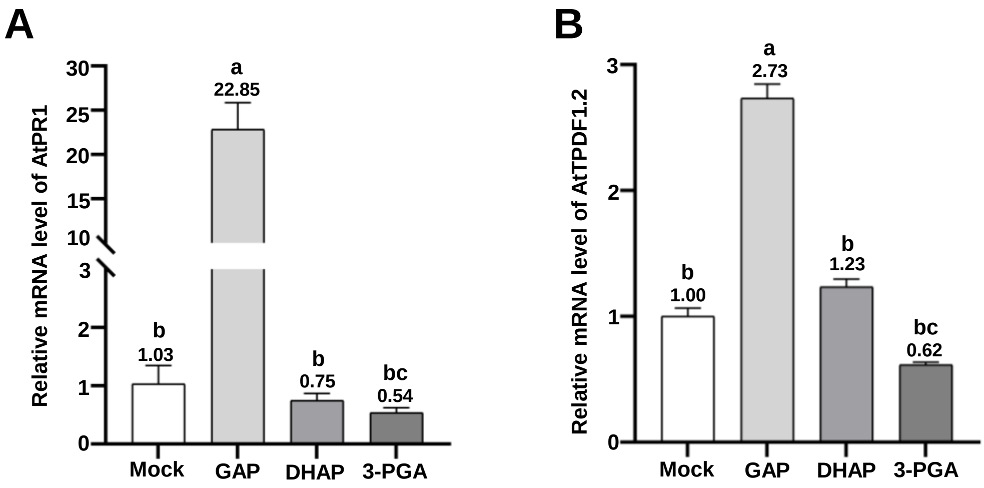


**Fig. S2. GAP specifically induces the expression of genes functioning in defense-related pathways in *Arabidopsis thaliana*.** RT-qPCR analysis of *AtPR1* (**A**) and *AtPDF1.2* (**B**) expression levels which were treated with 1mM GAP, DHAP, 3-PGA, or mock on *Arabidopsis* at 24 hrs. Error bars mean SD, and letters above the columns represent statistically significant differences (ANOVA, P<0.05).

**Data Set S1**.

Raw RNA-seq data obtained from GAP-treated *N. benthamiana* leaves related to Figure 5.

**Table S1 Sequence of primers used in this study**

| Primer | Sequence (5' to 3') | Experiment |
|---|---|---|
| AtTPT(KpnI)-F | GGGGTACCATGGAGTCACGCGTGCTG | pMDC32-AtTPT-3Flag for transgene |
| AtTPT(SpeI-TAG)-R | GGACTAGTTGCTTTCTTTCCTTGCCGTTTC | |
| AtTPT(H184A)-F | TGTCACGCCTTAGGCGCTGTCACTAGCAATGTC | Alanine substitution mutagenesis of AtTPT for pMDC32-AtTPT$^{H184A}$-3Flag construction |
| AtTPT(H184A)-R | GACTGCGACTGGTATCAATACCTTGAGGAGGTT | |
| AtTPT(R360A)-F | GGAAACGTTCTGAAAGCTGTGTTCGTGATCGGT | Alanine substitution mutagenesis of AtTPT for pMDC32-AtTPT$^{R360A}$-3Flag construction |
| AtTPT(R360A)-R | AACCGCGTGAGTCAGCGGTGCAACCCTCTCCAA | |
| qP0$^{Br}$-F | TCAGCTTCCTCTCCTTCTCG | BrYV RNAs RT-qPCR |
| qP0$^{Br}$-R | ATTGCTCGCAAGAGATCGTT | |
| qAtTPT-F | CTATGGTTGTCTCTAGCTCCTG | AtTPT RT-qPCR |
| qAtTPT-R | GATGGAGATGTAAGCGTAGACA | |
| qNbEF1A-F | GATTAATGAGCCCAAGAGGCC | NbEF1A RT-q PCR |
| qNbEF1A-R | AGTTTCCACACGACCAACAGG | |
| qNbPR1-F | TTAGCAGCCGTCATGAAATCGT | NbPR1 RT-qPCR<br>(*Continued to next page*) |
| qNbPR1-R | GGCGTAGAACCTTTAACCTGGGA | |

| | | |
|---|---|---|
| qNbPR10-F | GAAGAAGAACACAATGAAGGCA | NbPR10 RT-qPCR |
| qNbPR10-R | CAGTAGGATTGGCAAGAAGGTA | |
| qNbPDF1.2-F | GGAAATGGCAAACTCCATGCG | NbPDF1.2 RT-qPCR |
| qNbPDF1.2-R | ATCCTTCGGTCAGACAAACG | |
| qNbSABP2-F | ATGCCATGGAGGTTGGAGTTG | NbSABP2 RT-qPCR |
| qNbSABP2-R | ATTCATACCACCAAGACTATG | |
| AtActin2-F | GCACCCTGTTCTTCTTACCG | AtActin2 RT-qPCR |
| AtActin2-R | AACCCTCGTAGATTGGCACA | |
| qAtPR1-F | TGGTCACTACACTCAAGTTGTT | AtPR1 RT-qPCR |
| qAtPR1-R | GCTTCTCGTTCACATAATTCCC | |
| qAtPR2-F | CGTTGTGGCTCTTTACAAACAA | AtPR2 RT-qPCR |
| qAtPR2-R | AGCTCTGAACGTTTTCTTGAAC | |
| qAtPR5-F | AGGATTTGAATTGACTCCAGGT | AtPR5 RT-qPCR |
| qAtPR5-R | CCATCGCCTACTAGAGTGAATT | |
| qAtPDF1.2-F | CTTATCTTCGCTGCTCTTGTTC | AtPDF1.2 RT-qPCR |
| qAtPDF1.2-R | TGGGAAGACATAGTTGCATGAT | (*Continued to next page*) |

| | | |
|---|---|---|
| qAtNPR1-F | ATGATTTCTACAGCGACGCTAA | AtNPR1 RT-qPCR |
| qAtNPR1-R | GACTTCGTAATCCTTGGCAATC | |
| qAtICS1-F | CAATTAGGTGTCTGCAGTGAAG | AtICS1 RT-qPCR |
| qAtICS1-R | TTCGGACTGGTTAGTAAGTCAC | |
| qAtMYC2-F | CGGATCAGGAGTACAGGAAAAA | AtMYC2 RT-qPCR |
| qAtMYC2-R | GAAAAACCATTCCGTATCCGTC | |
| qAtCYP79B3-F | CAATCAAGAGGCTTATGTTCGG | AtCYP79B3 RT-qPCR |
| qAtCYP79B3-R | TAGATCCAATCCCGTAAGCATC | |
| qAtERF1-F | TTTCTCTACGGTCTAATCGAGC | AtERF1 RT-qPCR |
| qAtERF1-R | TGACTTTCTTGAGCTTACGGAT | |
| TPT-LP | GGTGTGAGTGAAGGAGACAGC | Primers for Homozygous individuals Verification |
| TPT-RP | AGAGGGAGCGAATCTGATCTC | |
| LB1.3 | ATTTTGCCGATTTCGGAAC | |
